# The spectral slope as a marker of excitation/inhibition ratio and cognitive functioning in multiple sclerosis

**DOI:** 10.1101/2023.01.23.525139

**Authors:** Fahimeh Akbarian, Chiara Rossi, Lars Costers, Marie B D’hooghe, Miguel D’haeseleer, Guy Nagels, Jeroen Van Schependom

**Affiliations:** Department of Electronics and Informatics (ETRO), Vrije Universiteit Brussel, Brussels, Belgium; AIMS lab, Vrije Universiteit Brussel, Center for Neurosciences, Brussels, Belgium; icometrix, Leuven, Belgium; National MS Center Melsbroek, Melsbroek, Belgium; UZ Brussel, Department of Neurology, Brussels, Belgium; St Edmund Hall, University of Oxford, Oxford, UK

**Keywords:** Aperiodic, 1/f spectral slope, aperiodic exponent, Excitation/inhibition balance, Magnetoencephalography, Resting state, Spectral power

## Abstract

**Background:** Multiple sclerosis (MS) is a neurodegenerative disease characterized by neuronal and synaptic loss, resulting in an imbalance of excitatory and inhibitory synaptic transmission and potentially cognitive impairment. Current methods for measuring the excitation/inhibition (E/I) ratio are mostly invasive, but recent research combining neurocomputational modeling with measurements of local field potentials has indicated that the slope with which the power spectrum of neuronal activity captured by electro- and/or magnetoencephalography rolls off, is a non-invasive biomarker of the E/I ratio. A steeper roll-off is associated with a stronger inhibition. This novel method can be applied to assess the E/I ratio in people with multiple sclerosis (pwMS), detect the effect of medication such as benzodiazepines, and explore its utility as a biomarker for cognition.

**Methods:** We recruited 44 healthy control subjects and 95 people with multiple sclerosis (pwMS) who underwent resting-state magnetoencephalographic recordings. The 1/f spectral slope of the neural power spectra was calculated for each subject and for each brain region.

**Results:** As expected, the spectral slope was significantly steeper in pwMS treated with benzodiazepines (BZDs) compared to pwMS not receiving BZDs (p = 0.01). In the sub-cohort of pwMS not treated with BZDs, we observed a steeper slope in cognitively impaired pwMS compared to cognitively preserved pwMS (p = 0.01) and healthy subjects (p = 0.02). Furthermore, we observed a significant correlation between 1/f spectral slope and verbal and spatial working memory functioning in the brain regions located in the prefrontal and parietal cortex.

**Conclusions:** In this study, we highlighted the value of the spectral slope in MS by quantifying the effect of benzodiazepines and by putting it forward as a potential biomarker of cognitive deficits in pwMS.

## 1. Introduction

Multiple sclerosis (MS) is an inflammatory and neurodegenerative disease of the central nervous system (1). Synaptic and neuronal loss are two pathological hallmarks in multiple sclerosis that lead to network dysfunction and disability and contribute to cognitive impairment (2–4). Despite the ongoing debate regarding which types of neurons and synapses are more likely to be affected by multiple sclerosis, the overall loss of synapses and neurons results in an imbalance of excitatory and inhibitory synaptic transmission (3,5). As a strict balance between excitatory and inhibitory synapses is a prerequisite for healthy brain function (6,7), the excitation/inhibition (E/I) ratio could be an important parameter to investigate in multiple sclerosis.

Current methods for measuring the E/I ratio are mainly invasive, such as single-unit (8) or voltage-clamp (9) recordings and can only be applied to relatively small population of cells, making them relatively non-applicable in a clinical setting or for in vivo population-level analyses. Animal models (10)and post-mortem tissue analysis (11) are two other approaches. While these approaches provide valuable insights, they have limitations regarding their applicability to clinical settings and the ability to capture real-time dynamics in living individuals. Magnetic resonance spectroscopy (MRS) (12) is another technique to estimate the E/I ratio by measuring the concentration of neurotransmitters across the brain. However, the temporal resolution of MRS is limited, and several minutes of recording (∼10 min) are needed to capture a single measure of neurotransmitter concentration (13).

Throughout the years, there has consistently been a demand for in vivo measurement techniques to assess the excitatory/inhibitory (E/I) ratio in a clinical context. Recent research combining neurocomputational modeling with measurements of local field potential has shown that the 1/f roll-off of power spectrum density of neuronal activity measured by electro- and/or magnetoencephalography (EEG/MEG) is an easily accessible biomarker of the E/I ratio (14). Further, a recent optogenetic study showed that the 1/f spectral slope captures fluctuations in the E/I balance (15), with a flatter slope being associated with a higher E/I ratio (14). This indicates that a steeper roll-off is associated with a stronger inhibition.

Several studies have corroborated this concept and have shown that the 1/f spectral slope decreases with age (16,17) and is modulated by task (18) and neuropsychiatric conditions like schizophrenia (19). Moreover, the relationship between the spectral slope and arousal and cognitive load has been revealed in several studies: Kozhemiako et al (20) demonstrated that the average spectral slope becomes progressively steeper during the transition from wakefulness to non-rapid eye movement (NREM) sleep and finally to rapid eye movement (REM) sleep. Similarly, Schneider et al. (21) showed that the spectral slope effectively differentiates between the different sleep stages, suggesting its potential as a sleep state marker. During these sleep stages, there is an increase in the steepness of the spectral slope relative to the awake state. Regarding the association between the 1/f spectral slope and cognitive load, it has been observed that in typically developing children, the spectral slope shows dynamic characteristics and becomes flatter during higher cognitive demanding tasks (22).

Presently, only a few studies have evaluated the spectral slope in neurological diseases. The 1/f slope has been found to be flatter in adolescents with Attention-deficit/hyperactivity disorder (ADHD) on a dual performance attention “stopping task” compared to healthy controls (23). Furthermore, the 1/f slope in resting state EEG was shown to be steeper in schizophrenia patients compared to healthy subjects (24). A steeper 1/f slope was also observed following dopaminergic treatment in patients with Parkinson’s disease (PD), highlighting the potential utility of the 1/f spectral slope to detect the non-invasive treatment response in patients with PD (25).

In this paper, we first compared the spectral slope between pwMS with and without benzodiazepine treatment. As benzodiazepines are known to enhance the inhibitory effect of γ-Aminobutyric acid (GABA) neurotransmitter (26), we expect a decrease in E/I ratio and, consequently, a steeper spectral slope. As benzodiazepines are also known to increase beta power (27,28), we will incorporate control measures in the analysis to account for any periodic beta modulation resulting from the benzodiazepine use.

Further, previous post-mortem (5,29) and rodent (29) studies in MS have suggested that inhibitory interneurons are more susceptible to demyelination, leading to increased loss of inhibitory synapses. As network disinhibition has been suggested to be associated with cognitive impairment (5), we hypothesize that a decrease in inhibition – and thus an increase in E/I ratio and a flatter spectral slope, will be accompanied by increased cognitive impairment in MS. Due to the subjective and challenging nature of cognitive assessment in multiple sclerosis within a clinical setting, the potential use of the 1/f spectral slope as a biomarker can be valuable in evaluating cognitive function.

## 2. Methods and materials

### 2.1. Participants

MEG data were acquired in 139 people with MS (pwMS) during resting-state eyes-closed; 23 pwMS treated with benzodiazepines (MS(BZD+)), 72 not treated (MS(BZD-)) and 44 healthy subjects aged between 18 and 65 years. All pwMS were recruited at the National MS Center Melsbroek and had to have been diagnosed with multiple sclerosis according to McDonald’s revised criteria (30) and had an Expanded Disability Status Scale (EDSS) score (31) of less than or equal to 6. A relapse or corticosteroid treatment within the preceding six weeks, as well as a pacemaker, dental wires, major psychiatric disorders, or epilepsy were all criteria for exclusion from the study. MS patients was consisted of 84.21% or 80 relapsing–remitting MS (RRMS), 7.3% or seven primary-progressive MS (PPMS), 6.3 % or six secondary progressive MS (SPMS) and 2.1% or two clinically isolated syndrome (CIS) patients. We provide a detailed description of the included cohorts of people with MS (pwMS) and healthy controls in Table 1.

**Table 1.** Description of subjects. We report the mean values and standard deviations from the different clinical parameters. For EDSS the median and interquartile range (IQR) is shown. The comparisons were performed using permutation testing with N = 5000 for all parameters except sex for which a chi-squared test was used. CIS, clinically isolated syndrome; HC, healthy control; MS(BZD-)_CI, MS(BZD-) cognitively impaired; EDSS, expanded disability status scale; MS(BZD-)_CP, MS(BZD-) cognitively preserved; PPMS, primary-progressive multiple sclerosis; RRMS, relapsing–remitting multiple sclerosis; SPMS, secondary progressive multiple sclerosis.

|  | HC | MS sub-groups |  | MS(BZD-) subgroups |  | Group differences (p-values) |  |  |  |
| --- | --- | --- | --- | --- | --- | --- | --- | --- | --- |
|  | HC | MS(BZD+) | MS(BZD-) | MS(BZD-)_CI | MS(BZD-)_CP | MS(BZD+) vs MS(BZD-) | HC vs MS(BZD-)_CI | HC vs MS(BZD-)_CP | MS(BZD-)_CI vs MS(BZD-)_CP |
| *N* | 44 | 23 | 72 | 16 | 56 |  |  |  |  |
| *Sex (M/F)* | 19/25 | 2/21 | 28/44 | 6/10 | 22/34 | **0.01** | 0.92 | 0.76 | 0.98 |
| *Age (yrs)* | 47±11 | 47±8 | 48±10 | 52 ±6 | 47±11 | 0.75 | 0.11 | 0.83 | 0.06 |
| *Education (yrs)* | 15.2±2 | 13±2 | 14±2 | 13±2 | 14±3 | 0.22 | **0.007** | **0.02** | 0.51 |
| *Disease duration(yrs)* | - | 14±6 | 15±10 | 14 ±9 | 16±9 | 0.42 | - | - | 0.57 |
| *EDSS median [IQR]* | - | 3.5[2.75-4.5] | 2.5[2-3.625] | 3.25 [2-4] | 2.5 [2-3.5] | **0.02** | - | - | 0.17 |
| *RRMS/PPMS /SPMS/ CIS* | - | 18/3/2/0 | 62/4/4/2 | 12/2/1/1 | 50/2/3/1 | - | - | - | - |
| *Cognitive assessment* |  |  |  |  |  |  |  |  |  |
| *SDMT* | 54 ±9 | 46±6 | 48±12 | 38±9 | 51±11 | 0.53 | **P < 0.001** | 0.13 | **0.0001** |
| *VGLT* | 66 ±7 | 63±9 | 62±11 | 48±10 | 66 ±8 | 0.62 | **P < 0.001** | 0.82 | **P < 0.001** |
| *COWAT* | 19±4 | 18±5 | 15±5 | 12±4 | 16±4 | 0.03 | **P < 0.001** | 0.052 | **0.004** |
| *BVMT* | 29±5 | 25 ±6 | 25±7 | 17±5 | 27±5 | 0.97 | **P < 0.001** | 0.29 | **P < 0.001** |

### 2.2. MEG data acquisition

The MEG data was collected at the CUB Hôpital Erasme (Brussels, Belgium) on an Elekta Neuromag Vectorview scanner (Elekta Oy, Helsinki, Finland) for the first 30 pwMS and 14 healthy subjects and the rest of subjects were scanned using an upgraded scanner, Elekta Neuromag Triux scanner (MEGIN, Croton Healthcare, Helsinki, Finland). Both MEG scanners used sensor layout as 102 triple sensors, each consisting of one magnetometer and two orthogonal planar gradiometers and were placed in a lightweight magnetically shielded room (MaxshieldTM, Elekta Oy, Helsinki, Finland).

The MEG data were acquired in the eyes-closed resting-state condition for 8 minutes. All participants were asked to stay still and close their eyes. MEG signals were recorded with a 0.1–330 Hz passband filter at 1 kHz sampling rate. Throughout the whole data acquisition process, the head positions of the subjects were tracked continuously using four head-tracking coils. The location of these coils and at least 400 head-surface points (on the nose, face, and scalp) with respect to anatomical fiducials were determined with an electromagnetic tracker (Fastrak, Polhemus, Colchester, Vermont, USA). An electrooculogram (EOG) and electrocardiogram (ECG) were simultaneously recorded with MEG signal acquisition.

### 2.3. MEG processing and parcellation

The preprocessing steps have been done using Oxford’s Software Library (OSL) pipeline built upon FSL, SPM12 (Welcome Trust Centre for Neuroimaging, University College London) and Fieldtrip. First, we used the temporal extension of the signal space separation algorithm (MaxfilterTM, Elekta Oy, Helsinki, Finland, version 2.2 with default parameters) to remove the external interferences and correct head movements (32). After applying of an FIR antialiasing lowpass filter, we downsampled the data to 250 Hz.

Using OSL’s RHINO (https://github.com/OHBA-analysis) algorithm, data were automatically coregistered with the subject’s T1 image: the head shape points were coregistered to the scalp extracted using FSL’s BETSURF and FLIRT (33,34), and then transformed into a common MNI152-space (35). Data were filtered between 0.1 and 70 Hz and then a 5^th^ order butterworth filter (between 49.5 and 50.5 Hz) was applied to remove the power line noise.

A semi-automated independent component analysis (ICA) algorithm was employed to visually identify and remove ocular and cardiac artefacts based on the correlation of the components’ time series with EOG and ECG signals respectively. Next, we used a linearly constrained minimum variance (LCMV) beamformer to project the MEG data to a source space (36–38). The source-reconstructed data were then parceled using a parcellation atlas including 42 parcels (39). The first principal component (PC) of the different time series within each parcel was used as the time series for that parcel. As previously discussed, the parcellation atlas covered the entire cortex and did not include subcortical areas (27,39).

### 2.4. Power spectral analysis and estimation of the aperiodic components

For each parcel, the power spectral density (PSD) was calculated using the default Welch function in SciPy with a window length of 500 samples (2 seconds). The power spectrum density consists of two distinct components: 1) an aperiodic component, reflecting 1/f roll-off characteristics, modeled with an exponential fit, 2) periodic components, reflecting putative oscillations (band-limited peaks) and modeled as Gaussians. We used the “fitting oscillations and one over f” (FOOOF) algorithm (40) to estimate the 1/f spectral slope. This algorithm aims to extract the periodic component by applying an iterative Gaussian fit to all oscillations. By modeling the oscillations and removing it from the fundamental power spectrum, an ideal 1/f slope can be extracted.

We defined the fitting frequency range between 20 Hz to 45 Hz to avoid the oscillatory peaks and to be in line with the methodology as presented by Gao et al (14), suggesting the stronger relationship between E/I ratio and spectral slope within the intermediate frequency range.

#### 2.4.1. Static analysis

For the static analysis, we used the first 2-minutes of recorded MEG data in order to reduce the possible effect that pwMS with BZD were more likely to fall asleep during the recording. For each subject, the power spectrum density of 42 brain parcels was calculated separately, and the 1/f spectral slopes were extracted using the FOOOF algorithm.

#### 2.4.2. Dynamic analysis

To perform a dynamic analysis, we used the entire 8 minutes of recorded MEG data, segmenting it into 16 sections of 30 seconds each. We then performed a static analysis within each 30-second data segment to evaluate the 1/f spectral slopes.

### 2.5. Estimation of the periodic power in the beta frequency range

First, we separated the periodic component by correcting the power spectrum density for the aperiodic component fitted within the frequency range of 13 to 45 Hz. The beta frequency range was then defined as between 13 to 30 Hz. To avoid the strong peak at 16.5 Hz, which was likely caused by construction works at the time of data acquisition at the hospital, we excluded power values at 16.5 Hz. To calculate the periodic beta power, the oscillatory beta power was averaged within the defined frequency band.

### 2.6. Neuropsychological assessment

Neuropsychological tests have been done on the day of the MEG recording for all subjects. The neuropsychological tests included the Symbol Digit Modalities Test (SDMT, (41)) to capture information processing speed, the Dutch version of the California Verbal Learning Test (CVLT-II, (42)), Dutch version: VGLT to assess verbal memory, the Controlled Oral Word Association Test (COWAT, (43)) to assess verbal fluency and the Brief Visuospatial Memory Test (Revised; BVMT-R, (44)) to assess spatial memory. We included the SDMT, VGLT and BVMT-R as they constitute the BICAMS test (45) and – in addition – acquired the COWAT to capture verbal fluency. Fatigue is assessed by the Fatigue Scale for Motor and Cognitive Function (FSMC, (46)) and depression by Beck’s depression inventory (BDI, (47)).

### 2.7. Cognitive impairment in multiple sclerosis

In accordance with the literature (48,49), impairment in each test of the BICAMS battery was defined as scoring 2 standard deviations below the mean normative values for each cognitive task, and overall cognitive impairment was defined as failing at all three BICAMS tests. We used a linear regression model similar to the model suggested by Costers et al (50), on the group of healthy subjects (matched in age and sex) to correct the raw cognitive scores for age, sex, and educational level.

### 2.8. Statistics

All statistical analyses used non-parametric measures, which do not rely on assumptions about the distribution of the data. We present the U-statistic and effect size for all pairwise comparisons performed with the Mann-Whitney U test. When calculating tests for multiple cortical parcels, the Benjamini-Yekutieli false discovery rate (FDR) (51) correction for multiple comparisons was applied to the p-values. For all p-values, the cutoff of 0.05 was used. The analysis of covariances was done using an ANCOVA model (52). Correlation analyses were controlled for the Expanded Disability Status Scale (EDSS) and disease duration variables using partial Spearman correlation. The effect size of all analyses will be reported.

### 2.9. Ethics

The research was approved by the University Hospital Brussels’s local ethics committees of the University Hospital Brussels (Commissie Medische Ethiek UZ Brussel, B.U.N. 143201423263, 2015/11) and the National MS Center Melsbroek (2015-02-12). All participants also provided written informed consent.

A schematic of the study pipeline is shown in Figure 1.

**Figure 1.**
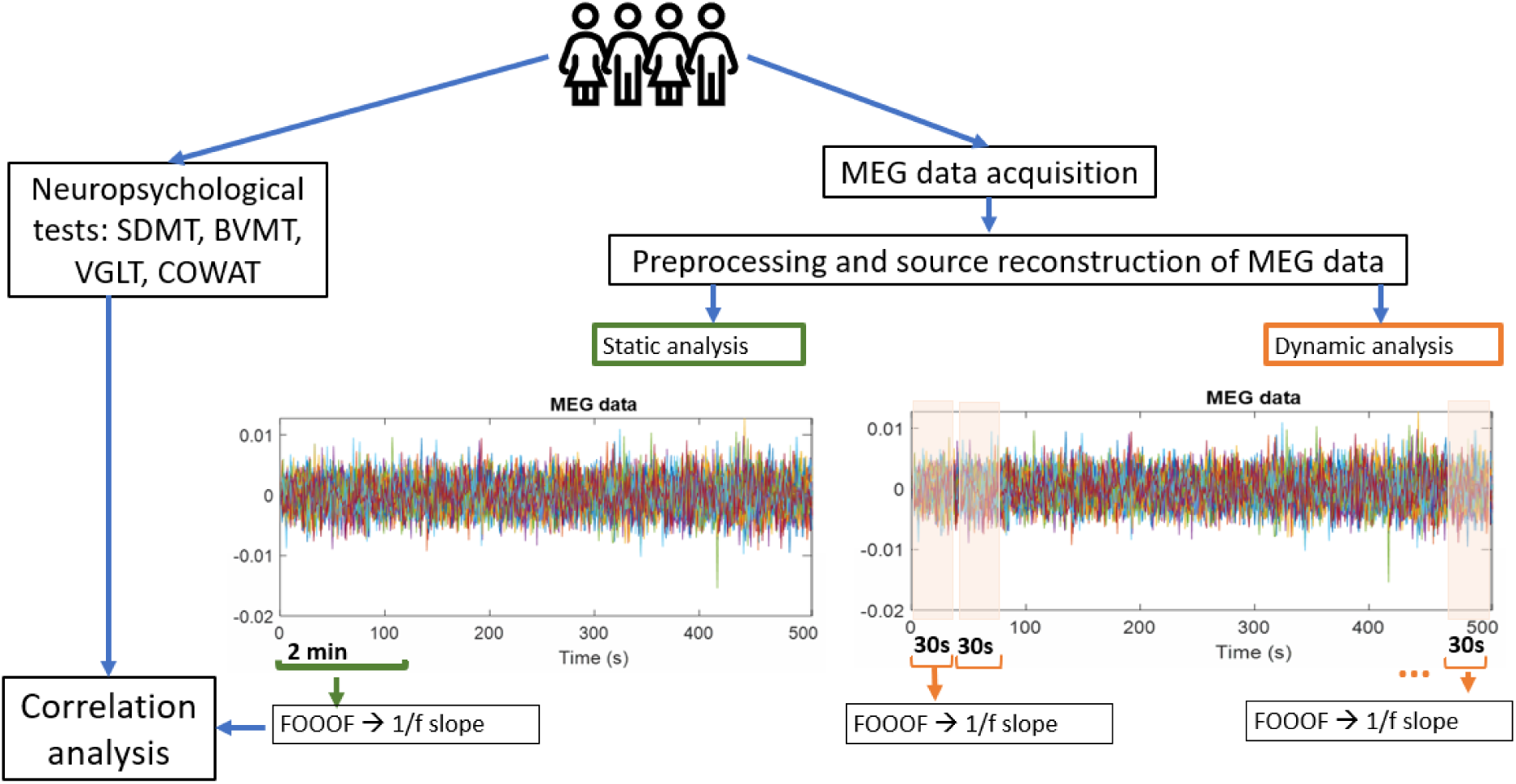
Pipeline of study. We obtained MEG resting-state eyes closed data and neuropsychological test results at the same day. The MEG data was processed to enable the extraction of the 1/f slope. This was then compared between people with and without benzodiazepines and later also correlated with cognitive test results.

## 3. Results

### 3.1. Static analysis of comparing MS populations: MS(BZD+) vs MS(BZD-)

#### 3.1.1. Whole brain

We present a whole-brain analysis by averaging the 1/f slope between 20 and 45 Hz across the 42 parcels for each subject. Next, we compared the averaged 1/f slopes between two groups of pwMS (MS(BZD+) vs MS(BZD-)). As expected, and shown in Figure 2, we observed a significant difference in 1/f slope in which pwMS treated with benzodiazepines having a steeper slope (p-value = 0.014, Mann-Whitney-U = 547, η^2^= 0.063, Cohen’s d = 0.51).

**Figure 2.**
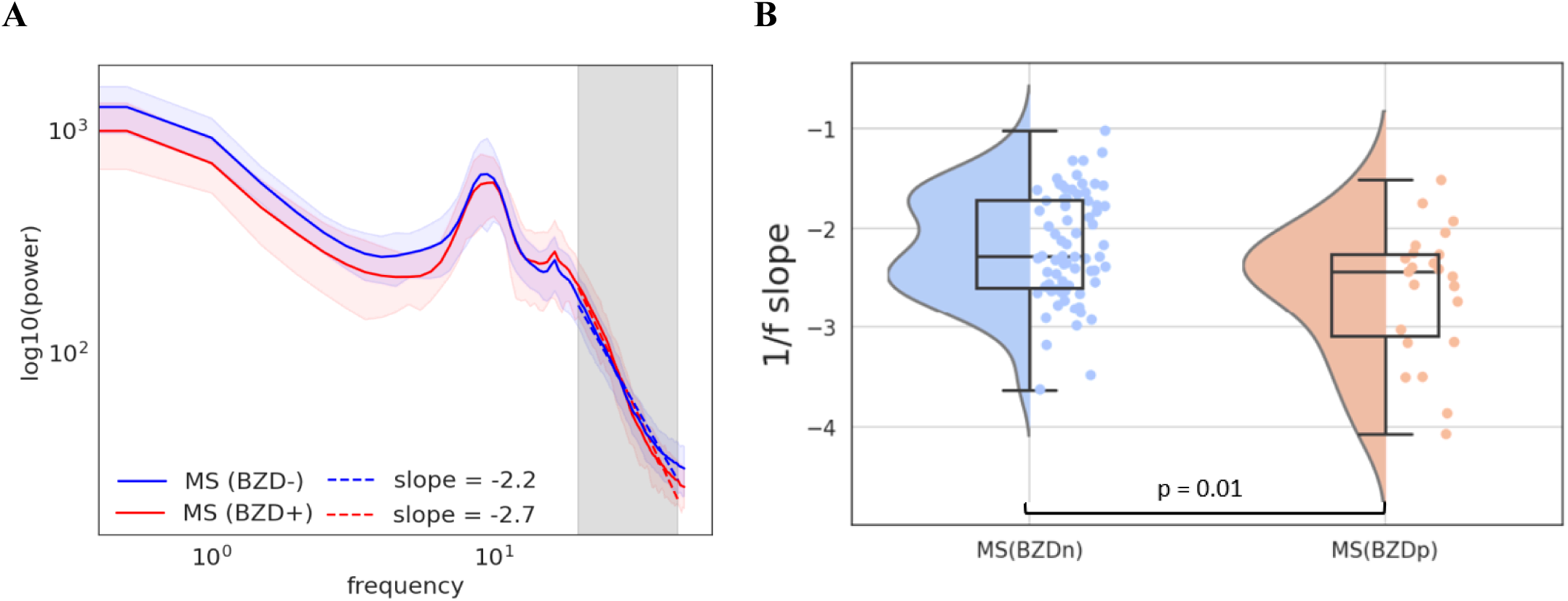
A) The spectra of both cohorts on a log-log scale and the corresponding 1/f slopes. The solid lines present the median and the shaded areas the median absolute deviation. The grey box indicates the area in which the roll-off was calculated [20-45 Hz]. **B)** Distribution of averaged 1/f slope across the whole brain. Each dot represents one subject. p-value = 0.014, Mann-Whitney-U = 1109, η2 = 0.063, Cohen’s d = 0.51.

#### 3.1.2. Parcel level

Next, we extracted the 1/f slope for each parcel separately and compared them between MS(BZD+) and MS(BZD-). A significant difference in 1/f slope was observed in 26 brain parcels after correction for multiple comparisons (51). The MS(BZD+) had a steeper slope compared to MS(BZD-). As it is shown in Figure 3, pwMS treated with benzodiazepines had steeper slopes in all brain parcels.

**Figure 3.**
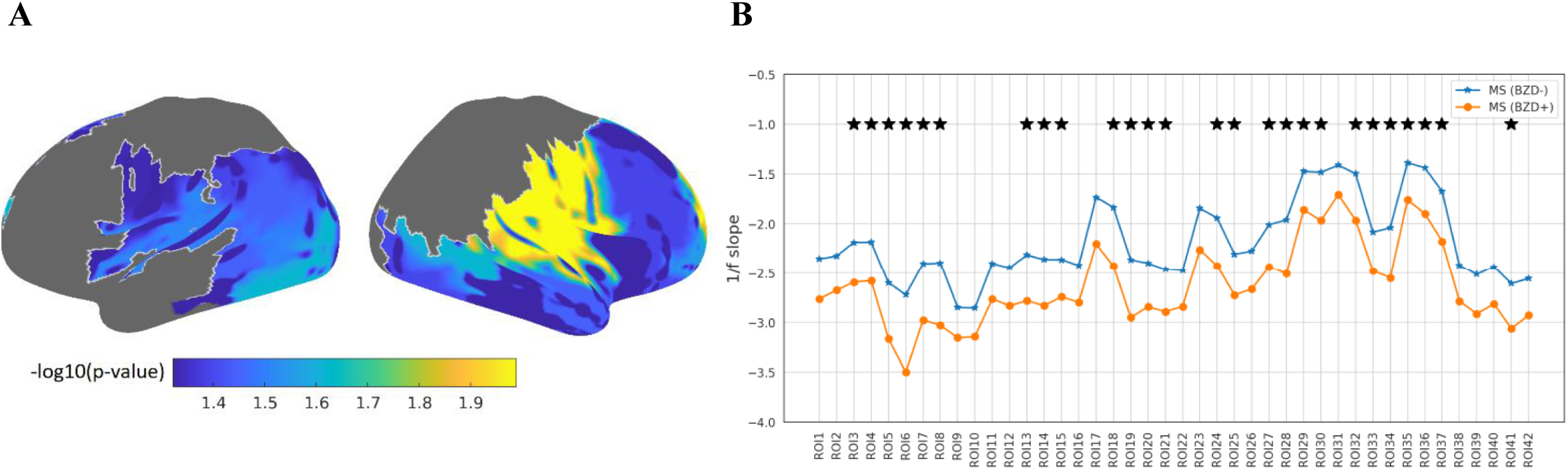
A) Comparison of slope in the population of pwMS who have received benzodiazepines (MS(BZD+) and pwMS who have not received benzodiazepines (MS(BZD-). MS(BZD+) has steeper slopes compared with MS(BZD-) when comparing across brain parcels. **B)** Slopes were steeper in MS(BZD+) for all brain parcels. The regions of interest (ROIs) are listed in Table S1. *p < 0.05 or -log10(p-value) > 1.3: statistically significant after false discovery rate correction for multiple comparison.

### 3.2. Dynamic analysis of comparing MS populations: MS(BZD+) vs MS(BZD-)

Within each 30-second segment of the data, we compared the 1/f slope of the pwMS that were treated with benzodiazepines and those that were not treated with them. The results of this dynamic analysis demonstrate that the difference between these two groups remains significant over the complete recording session (Supplementary materials. Figure S1).

### 3.3. Spectral slope and beta activity

To rule out the variations in periodic beta activity as a potential explanation for the observed group differences in spectral slope, we estimated the spectral power in the beta band. Figure 4 shows the cortical distributions of average periodic beta power in the two groups of pwMS (MS(BZD+) and MS(BZD-)).

**Figure 4.**
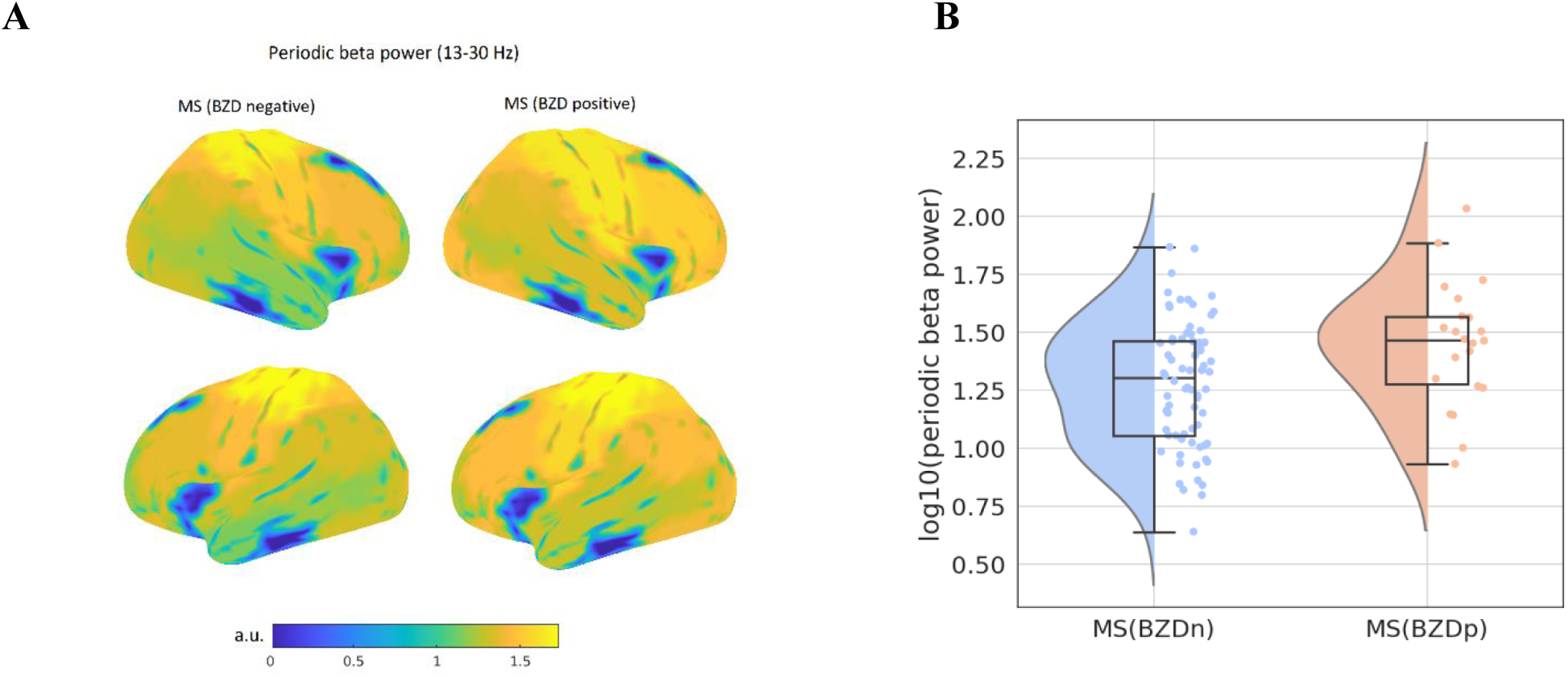
**A**) Cortical distributions of periodic beta power (13-30 Hz). Distributions are shown in the two groups of pwMS. Top: right hemisphere, bottom: left hemisphere. a.u., arbitrary units. **B)** Distribution of mean periodic beta power in each group of pwMS. Each dot represents one subject. (p-value = 0.041, Mann-Whitney-U = 593.0, η^2^ = 0.04, Cohen’s d = 0.42).

We observed that in both cohorts an increase in the periodic beta power corresponds to an increase in the negative slope of the spectra (r = -0.65, p < 0.001). Since there was a significant difference in average periodic beta power between the two groups (p-value = 0.041, Mann-Whitney-U = 593.0, η^2^ = 0.04, Cohen’s d = 0.42), we reran the comparison of 1/f slope between two groups using the ANCOVA model to control for periodic beta power. The difference in 1/f slope between two groups of pwMS (MS(BZD+) and MS(BZD-)) remained statistically significant (p-value = 0.02). Although the average periodic beta power difference over the entire brain significantly differed between the two groups, there was no significant difference at the parcel level after false discovery rate (FDR) correction.

### 3.4. Spectral slope and cognitive impairment

In the second part of this study, we investigated the relationship between 1/f slope as an index of cognitive functioning in the group of MS(BZD-) patients. Based on the criteria defined above, we divided the MS(BZD-) into two groups (n =72): cognitively impaired (n =16) and cognitively preserved (n = 56). We then evaluated between-group differences in averaged - across the whole brain- 1/f slope between three different groups: cognitively impaired pwMS, cognitively preserved pwMS, and healthy subjects. These three groups were matched in sex and age.

We observed a significant difference in slope between cognitively impaired pwMS and cognitively preserved pwMS (p-value = 0.013, Mann-Whitney-U = 262.0, η^2^= 0.08, Cohen’s d = 0.60). Average slopes were significantly steeper in the cognitively impaired pwMS compared to cognitively preserved pwMS. Similarly, we observed a significant difference in slope between cognitively impaired pwMS and healthy subjects (p-value = 0.029, Mann-Whitney-U = 482.0, η^2^= 0.07, Cohen’s d = 0.58). Average slopes were significantly steeper in the cognitively impaired pwMS compared to healthy subjects. Figure 5 shows the distribution of averaged 1/f slope across the whole brain for three groups.

**Figure 5.**
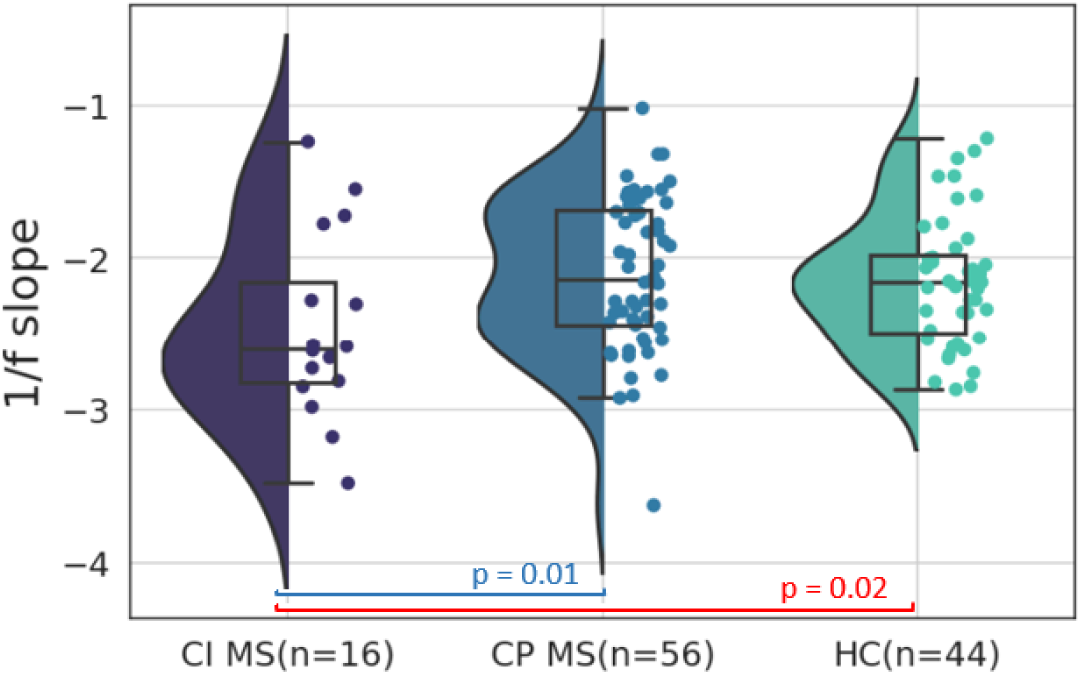
Distribution of averaged 1/f slope across the whole brain for three groups. Each dot represents one subject. CI MS: cognitively impaired pwMS. CP MS: cognitively preserved pwMS. HC: healthy control subjects.

We did not observe a significant difference in slope between cognitively preserved pwMS and healthy subjects (p-value = 0.50, Mann-Whitney-U = 1115.0, η^2^ = 0.005, Cohen’s d = 0.13).

Given the notable disparity in educational level among healthy individuals and cognitively impaired pwMS and cognitively preserved pwMS, we took into account the influence of education level on the findings. After controlling for education level, the results remained the same, with a p-value of 0.03 and a p-value of 0.69, respectively.

Furthermore, we also performed a correction for age, which was borderline significantly different (p-value = 0.06, see Table 1), in the comparison between cognitively impaired pwMS and cognitively preserved pwMS. After the correction was applied through the ANCOVA model, the results remained significant with a p-value of 0.04.

### 3.5. Correlation analysis between the 1/f slope and cognitive scores

To further explore the relationship between the 1/f slope and each cognitive score separately, we performed correlation analyses. We observed a significant correlation between the 1/f slope – averaged across the full brain – and BVMT (r = 0.26, p = 0.02) and VGLT (r = 0.24, p = 0.03) scores. A steeper slope was associated with lower BVMT and VGLT scores, as it is shown in Figure 6. There was no link between the 1/f slope and the SDMT or COWAT, neither in the whole brain nor in the individual parcels. All correlations were corrected for EDSS and disease duration. In general, the correlation between spectral slope and BVMT and VGLT was more significant in the brain parcels located in the prefrontal and parietal regions.

**Figure 6.**
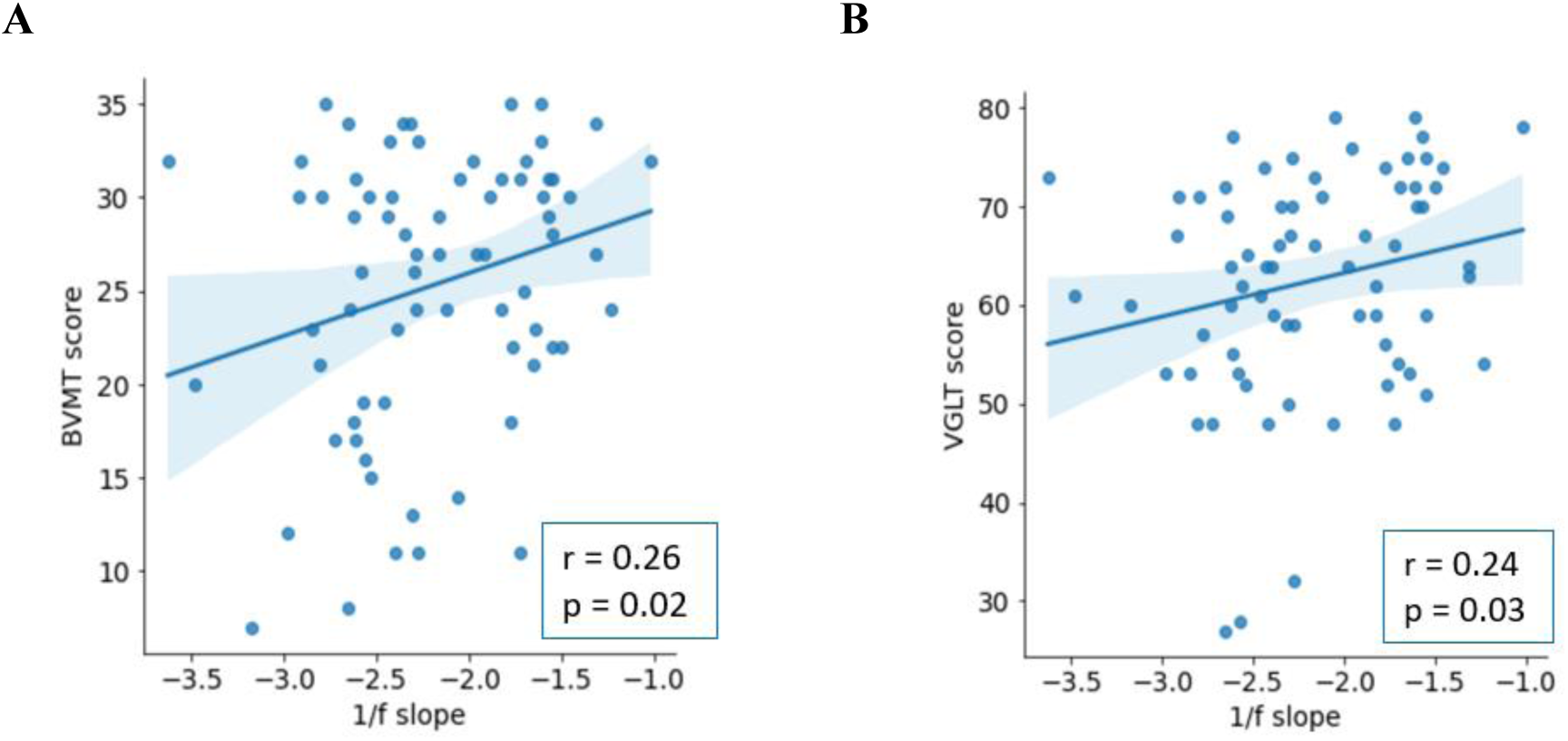
Correlation analysis between 1/f slope and BVMT scores (**A**) and VGLT scores (**B**). All correlations were calculated on the cohort of pwMS who have not treated with benzodiazepine: MS(BZD-).

We repeated the correlation analysis after excluding three pwMS with the lowest VGLT scores (outliers). We did not observe a significant correlation between the 1/f slope – averaged across the full brain – and VGLT (r = 0.22, p = 0.07) scores. However, the slope was still significantly correlated in the brain parcels located in the prefrontal and parietal regions.

The correlation analysis was repeated for the group of pwMS who had received benzodiazepine treatment (n = 23). We did not observe a correlation between 1/f slope and cognitive scores.

## 4. Discussion

### 4.1. The effect of benzodiazepines on 1/f slope in pwMS

In our study, we demonstrate that the 1/f spectral slope can distinguish between people with multiple sclerosis (pwMS) who receive benzodiazepines treatment and those who do not. Whereas our study did not include other markers of excitation/inhibition ratio (e.g., PET imaging), our results corroborate the idea that the 1/f slope can be interpreted as the E/I ratio in MS. Indeed, administration of BZDs resulted in a steeper slope which is typically interpreted as an increased inhibition. Future research should explore this idea, e.g., through an intrasubject design. The significant difference in 1/f slope was observed in large parts of the brain, namely the occipital, temporal, and prefrontal brain regions. In addition, while we focused on the first 2 minutes of data to reduce the potential effects of people falling asleep, we also showed that this effect can be consistently observed across the full recording.

The impact of benzodiazepines on beta power is widely recognized and documented (e.g., (28)). Consequently, an essential question arises: could the observed differences in slope be merely an epiphenomenon resulting from variances in beta power between the MS(BZD+) and MS(BZD-) groups? To address this concern, we accounted for periodic beta power as a covariate in our analysis. Even after considering this factor, the disparity in the 1/f slope remained statistically significant (p = 0.03) between the two sub-groups of pwMS. This finding indicates that the spectral slope serves as an independent metric that holds distinct informative value alongside beta periodic power.

### 4.2. Fitting frequency range selection

It is not trivial to choose the fitting frequency window and FOOOF modeling parameters. Therefore, we repeated the analyses for multiple fitting frequency ranges: 1) a broad-band frequency range (3-45 Hz), lower-border of the fitting range started from 3 Hz to avoid the impact of these low-frequency oscillations (53), 2) a frequency window including the beta peak (13- 45 Hz) and 3) a frequency window excluding the beta peak (30- 45 Hz) (53). The PSDs were not linear in log-log space when the broad-band frequency range and the frequency range between 13 and 45 Hz were considered. Therefore, we calculated two models with and without an additional knee and we observed a smaller value of the Akiake Information Criterion (54) for the models with a knee in the broadband frequency. These additional analyses demonstrated that the specific choice of frequency window did not affect our findings, see Supplementary Table S2.

### 4.3. Spectral slope and cognitive impairment

As neuronal and synaptic loss are well-known features of MS, we further investigated the association between the 1/f slope and cognitive functioning in the MS(BZD-) group. We focused on the MS(BZD-) group to avoid potential interference with the use of benzodiazepines in the analyses. We observed a significant correlation between 1/f spectral slope and verbal and spatial working memory, which was strongest in the regions involved in verbal and spatial memory, i.e., parietal (55) and prefrontal (56) regions. Although we expected flatter spectral slopes (typically interpreted as less inhibition) to be associated with worse cognitive functioning, we observed steeper slopes. Our results, on the contrary, suggest that a decrease in excitation or an increase in inhibition is associated with poorer cognitive performance.

Importantly, two clinical EEG studies evaluating aperiodic parameters in patients with schizophrenia and children with Attention-deficit/hyperactivity disorder (ADHD) have yielded similar findings (57,58). In addition, although schizophrenia is associated with decreased GABAergic inhibition in the cortex (59), Peterson et al found steeper 1/f slopes in schizophrenia patients compared to healthy control subjects, suggesting a compensatory increase in GABAergic activity. Our findings may indicate that a decrease in inhibitory interneurons may be overcompensated (3). It is also possible that there is a range of 1/f slopes that are considered to be cognitively normal whereas both slopes that are too steep or too flat can indicate cognitive impairment (58).

An alternative interpretation is that the 1/f spectral slope as a marker of the E/I ratio may not hold in these datasets. Baranauskas et al previously related the 1/f slope with the rate of transitions between cortical up and down states (60). In this interpretation, cognitive impairment would be associated with an abrupt transition between up and down states. Finally, Muthukumaraswamy and Lily interpreted the 1/f scaling as a side-effect of the dampening of harmonic oscillators (61). In that case, cognitively impaired individuals with MS may exhibit a weaker dampening of the alpha wave.

## 5. Study limitations

While we did not have benzodiazepine dosage information or treatment duration the obtained results are clearly significant. It could be expected that assessing these effects in a larger group of people treated with benzodiazepines may further elucidate if the observed effects are mainly due to a specific benzodiazepine or rather reflect an effect that is shared among the different BZD treatments.

Another potential limitation of this study was the fact that most of the pwMS who received BZDs were female (21 vs 2). Therefore, we compared the 1/f slope of the 19 male and 25 female healthy control subjects matched in age (p-value = 0.48) to rule out the effect of sex on the 1/f slope. We did not observe a significant difference in 1/f spectral slope between male and female healthy subjects (p-value = 0.76, Mann-Whitney-U = 224, η^2^= 0.002, Cohen’s d = 0.09). The pwMS who received BZDs also had a slightly higher Expanded Disability Status Scale (EDSS) score. Yet, including the EDSS in the ANCOVA model did not alter the results.

A final limitation is the acquisition of MEG data on two (slightly) different scanners. To investigate the effect of different scanners, we compared the whole brain averaged 1/f slope between 13 healthy subjects and 31 healthy subjects scanned by the first and the second scanner respectively. We did not observe a significant difference in 1/f spectral slope between two healthy groups of the first scanner and second scanner (p-value = 0.27, Mann-Whitney-U = 158, η^2^= 0.02, Cohen’s d = 0.34).

## 6. Conclusion

Multiple sclerosis (MS) is a neurological disorder characterized by progressive neuronal and synaptic degeneration, resulting in an imbalance in the transmission of excitatory and inhibitory signals. This disruption in the balance between excitatory and inhibitory synaptic transmission can potentially result in cognitive impairment in individuals affected by MS. While EEG and MEG studies have traditionally focused on examining group differences in oscillatory dynamics, recent research (14) underlines the importance of the 1/f slope as a potential biomarker for the excitation/inhibition ratio. In this study, we present the first investigation to highlight the utility of 1/f spectral slope to quantify the inhibitory effect of benzodiazepine usage in pwMS. We also demonstrated a correlation between the 1/f spectral slope and cognitive performance in pwMS who are not using benzodiazepines.

## Acknowledgment

We would like to thank all participants for their time, enthusiasm, and commitment to participate.

## Data availability

Data is available upon reasonable request to the corresponding author.

