## Supplementary materials for "The spectral slope as a marker of excitation/inhibition ratio and cognitive functioning in multiple sclerosis"

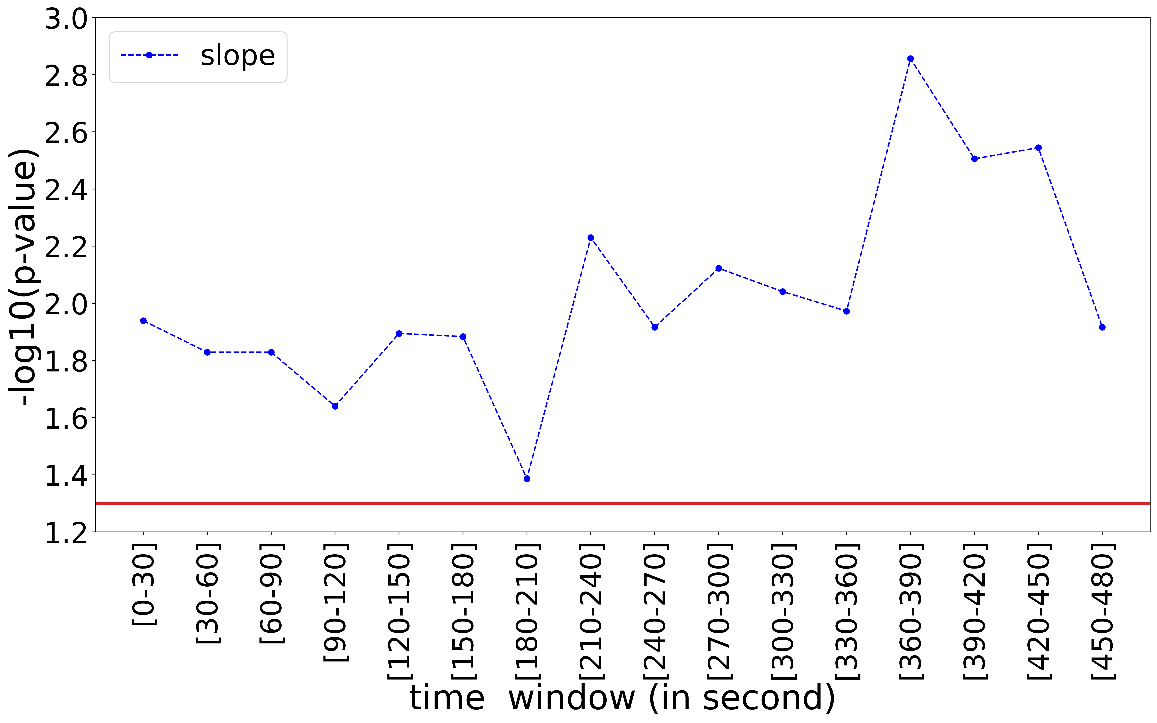

**Figure S1. Dynamic analysis.** On the y-axis, the p-values of the comparison of 1/f slope between MS(BZD+) and MS(BZD-) for different piece of data. One can clearly appreciate the difference being present in every piece data. Horizontal red line indicates the cut-off value of 1.3. The values of -log10(p-value) above 1.3 correspond to significant results.

**Table S1.** List of regions used in the parcel level analyses.

|  | ROI |
| --- | --- |
| 'ROI 1' | 'L Cuneus' |
| 'ROI 2' | 'R Cuneus' |
| 'ROI 3' | ' L Inf Occ' |
| 'ROI 4' | 'R Inf Occ' |
| 'ROI 5' | 'L Supramarginal' |
| 'ROI 6' | 'R Supramarginal' |
| 'ROI 7' | 'L Sup Temp' |
| 'ROI 8' | 'R Sup Temp' |
| 'ROI 9' | 'L Lat SMC' |
| 'ROI 10' | 'R Lat SMC' |
| 'ROI 11' | 'L Sup Parietal' |
| 'ROI 12' | 'R Sup Parietal' |
| 'ROI 13' | 'L Middle Occ' |
| 'ROI 14' | 'R Middle Occ' |
| 'ROI 15' | 'L Sup Occ' |
| 'ROI 16' | 'R Sup Occ' |
| 'ROI 17' | 'L Ant Temp' |
| 'ROI 18' | 'R Ant Temp' |
| 'ROI 19' | 'L Medial SMC' |
| 'ROI 20' | 'R Medial SMC' |
| 'ROI 21' | 'L Angular' |
| 'ROI 22' | 'R Angular' |
| 'ROI 23' | 'L VL PFC' |
| 'ROI 24' | 'R VL PFC' |
| 'ROI 25' | 'L Occ pole' |
| 'ROI 26' | 'R Occ pole' |
| 'ROI 27' | 'L Sup PFC' |
| 'ROI 28' | 'R Sup PFC' |
| 'ROI 29' | 'L Sup Dorsal PFC' |
| 'ROI 30' | 'R Sup Dorsal PFC' |
| 'ROI 31' | 'L Orbitofrontal' |
| 'ROI 32' | 'R Orbitofrontal' |
| 'ROI 33' | 'L Post Temp' |
| 'ROI 34' | 'R Post Temp' |
| 'ROI 35' | 'L Inf Dorsal PFC' |
| 'ROI 36' | 'R Inf Dorsal PFC' |
| 'ROI 37' | 'Medial PFC' |
| 'ROI 38' | 'Post Precuneus' |
| 'ROI 39' | ' Posterior Cingulate Cortex ' |
| 'ROI 40' | 'Ant Precuneus' |
| 'ROI 41' | 'L Inf Parietal' |
| 'ROI42' | 'R Inf Parietal' |

| Fitting frequency range | Fitting mode | Goodness of fit parameters: mean(std) | Goodness of fit parameters  (Each dot represents one subject) | 1/f aperiodic MS(BZDn) vs MS (BZDp) |
| --- | --- | --- | --- | --- |
| 3-45 Hz | **Knee** | R^2^ = 0.89 (0.03)  Error= 0.09 (0.01) | 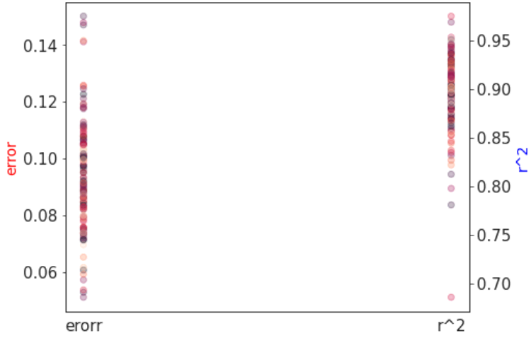 | p-value= 0.002  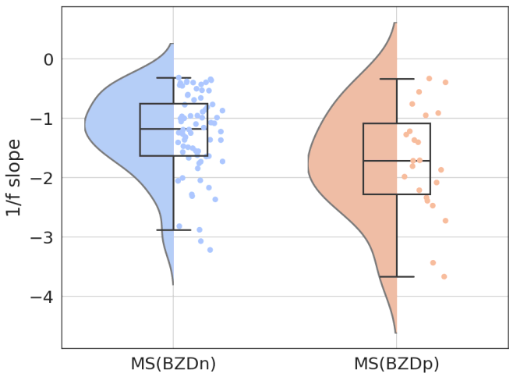 |
| 13-45 Hz | **Knee** | R^2^ = 0.93 (0.03)  Error= 0.06 (0.01) | 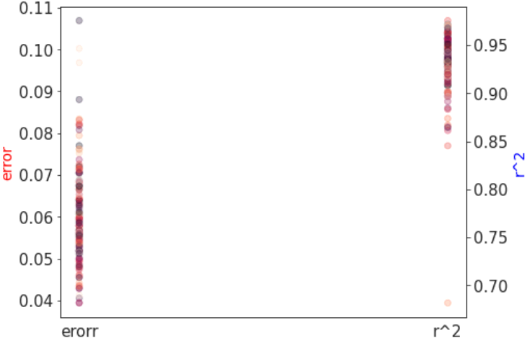 | 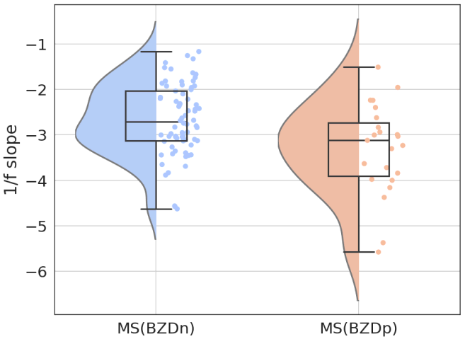p-value = 0.007 |
| 30-45 Hz | Fixed | R^2^ = 0.76 (0.10)  Error = 0.03 (0.003) | 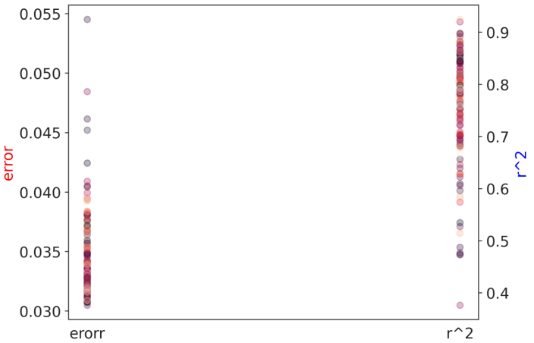 | 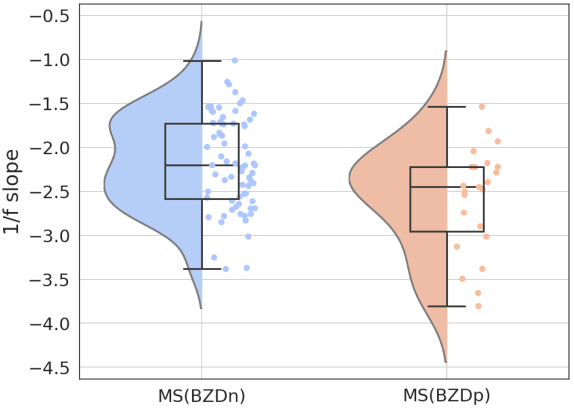p = 0.02 |
| 20-45 Hz | Fixed | R^2^ = 0.91 (0.04)  Error = 0.04 (0.008) | 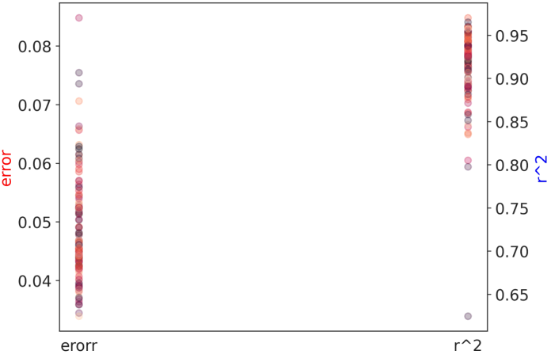 | p-value = 0.01  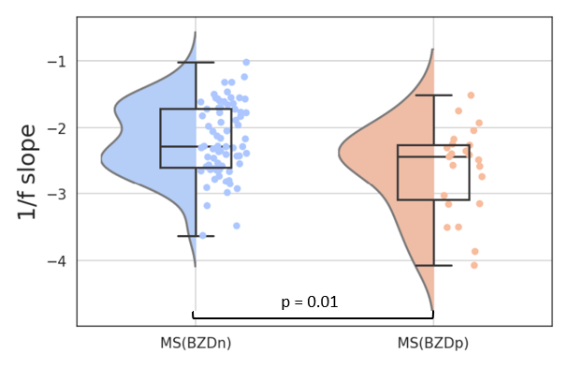 |

**Table S2.** Results of multiple fitting frequency range analysis
